# Uncovering mechanisms of global ocean change effects on the Dungeness crab (*Cancer magister*) through metabolomics analysis

**DOI:** 10.1101/574798

**Authors:** Shelly A. Trigg, Paul McElhany, Michael Maher, Danielle Perez, D. Shallin Busch, Krista M. Nichols

**Affiliations:** Division of Biological Sciences, Cell and Developmental Biology Section, University of California San Diego, La Jolla, California, USA; Genomic Analysis Laboratory, The Salk Institute for Biological Studies, La Jolla, California, USA; Conservation Biology Division, Northwest Fisheries Science Center, National Marine Fisheries Service, National Oceanic and Atmospheric Administration, Seattle, Washington, USA; Ocean Acidification Program, Office of Oceanic and Atmospheric Research and Northwest Fisheries Science Center, National Marine Fisheries Service, National Oceanic and Atmospheric Administration, Seattle, Washington, USA; Present address: School of Aquatic and Fishery Sciences, University of Washington, Seattle, Washington, USA

## Abstract

The Dungeness crab is an economically and ecologically important species distributed along the North American Pacific coast. To predict how Dungeness crab may physiologically respond to future global ocean change on a molecular level, we performed untargeted metabolomic approaches on individual Dungeness crab juveniles reared in treatments that mimicked current and projected future pH and dissolved oxygen conditions. We found 94 metabolites and 127 lipids responded in a condition-specific manner, with a greater number of known compounds more strongly responding to low oxygen than low pH exposure. Pathway analysis of these compounds revealed that juveniles may respond to low oxygen through evolutionarily conserved processes including downregulating glutathione biosynthesis and upregulating glycogen storage, and may respond to low pH by increasing ATP production. Most interestingly, we found that the response of juveniles to combined low pH and low oxygen exposure was most similar to the low oxygen exposure response, indicating low oxygen may drive the physiology of juvenile crabs more than pH. Our study elucidates metabolic dynamics that expand our overall understanding of how the species might respond to future ocean conditions and provides a comprehensive dataset that could be used in future ocean acidification response studies.

## INTRO

The continued increase in anthropogenic carbon dioxide emissions is leading to ocean acidification, with the ocean absorbing an average of 25% of human-caused emissions annually^1^. If atmospheric carbon dioxide concentration continues to rise at the current rate, the pH of oceans is predicted to fall 0.3-0.4 units by the end of the century^2,3^. This pH drop could exacerbate conditions in the U.S. Pacific Northwest, where the ocean pH is lower than that of the global ocean due to natural oceanographic processes including regional upwelling, and could pose a greater challenge to the marine inhabitants already coping with this lower pH. An additional and compounding factor to ocean acidification is ocean deoxygenation, which co-varies with pH and temperature. Given that global ocean temperatures will rise with global warming from continued greenhouse gas emissions, hypoxic zones are expected to increase in duration, intensity, and frequency^4^. It is not certain how future ocean acidification and deoxygenation environmental stress might affect important Pacific Northwest fisheries like the Dungeness crab fishery, which is the most lucrative and valued at more than $200 million annually^5^.

Ocean acidification is predicted to have negative indirect effects on Dungeness crab through loss of prey directly affected by ocean acidification^6^, but knowledge of how ocean acidification might directly impact Dungeness crab is limited. It has been postulated that Dungeness crab may exhibit limited tolerance for acid-base disturbances given past observations of Dungeness crab as weak osmoregulators^7–10^ and that acid-base balance and osmoregulation are tightly coupled in decapod crustaceans^11^. However, it was shown that following a brief two-week exposure to a future-predicted seawater pH of 7.4, adult Dungeness crab are able to acclimate by increasing hemolymph ion levels (bicarbonate, calcium, chloride, sulfate, and sodium), and by decreasing both oxygen consumption and nitrogen excretion^12^. On the other hand, Dungeness crab larvae have shown reduced survival and development rate in response to a 45-day exposure to the projected future seawater pH of 7.5 and 7.1^13^. Moreover, adult Dungeness crab have shown behavioral responses to declining oxygen conditions (21-1.5 kPa pO_2_ over a 5-hour period), including reduced feeding and a preference for the area with the highest pO_2_ level when placed in a seawater oxygen gradient (2.5-10.5 kPa pO_2_ for 1 hour)^14^. Dungeness crab also have shown physiological responses to declining oxygen conditions (18-3 kPa pO2 over a 6-hour period), including redistributing hemolymph to high-energy-demand tissues^15^. Despite these clear biological responses to low pH and oxygen, the biochemical mechanisms underlying Dungeness crab response to combined pH and oxygen stress have not yet been defined.

One way to gain broad insight into the biochemical processes underlying the physiological status of organisms is through surveying metabolomes, which is now possible for nearly any species with the advancement of high-throughput metabolite profiling, also known as metabolomics. Exploratory untargeted metabolomics approaches can offer unbiased analyses of the composition of all detectable metabolites for the rapid and quantitative detection of stress responses, which often leads to the development of targeted approaches and identification of stress-indicating biomarkers^16,17^. Functional analyses using pathway inference can subsequently be performed by integrating metabolomics data with databases and other “-omics” datasets using bioinformatics tools to ultimately establish causal networks between different experimental conditions and outcomes. Untargeted metabolomics in the context of understanding ocean acidification has recently been applied to reef-building coral, and revealed metabolite profiles that were predictive of primary production activity and molecules that could be used as potential biomarkers of ocean acidification^18^.

To better understand which biochemical pathways might be altered in the response of Dungeness crab to ocean acidification, we applied untargeted metabolomics and lipidomics to individual Puget Sound juveniles exposed to current pH (7.85) and future pH (7.45) conditions for an average of 32 days. To account for the dissolved oxygen (DO) levels that naturally co-vary with pH, we included both ambient oxygen (8.9 mg/L or saturated O_2_) and low oxygen (3.0 mg/L or 33% O_2_ saturation) treatments with our pH conditions in a factorial design to understand how pH, oxygen, and/or their interaction might influence metabolite abundances.

## RESULTS

### Juvenile Dungeness crab general metabolome and lipidome composition

Untargeted lipidomics and metabolomics were carried out on 60 individual juvenile crabs exposed to ocean acidification treatments (**Table 1** and **Supplementary Table 1**) through the entire duration of their first juvenile instar until 2 days after molting to their second juvenile instar (**Supplementary Table 2**). From the lipidomics and metabolomics profiles generated by the West Coast Metabolomics Core using in-house open-access bioinformatics pipelines^19,20^ (see Methods for additional details), a total of 3113 lipids and 651 general metabolites were detected of which 88% (3320/3764 total detected compounds) had spectra that did not match LipidBlast^21^ or BinBase^22^ records and were classified as unknown compounds (**Supplementary Table 3** and **4**). Eighteen lipids were detected in fewer than 50% of individual profiles and were therefore excluded from all downstream analyses. The MS identification techniques found 29/281 metabolite classes among the 246 compounds detected and listed in Human Metabolome Database (HMDB)^23^ and 13/77 lipid classes among 195 compounds detected and listed in LIPID MAPS Structural Database^24^ (**Supplementary Fig. 1**). Among the largest classes represented in all identified compounds were LIPID MAPS classes glycerophosphocholines, triradylglycerols, and fatty acids, and HMDB classes organic acids, carboxylic acids, and organic oxygen compounds, consistent with previously observed *Cancer magister* body composition proportions of these compounds^25,26^.

**Table 1.** Summary of water chemistry for experimental treatments showing the mean ± standard deviation for each parameter. Temperature (temp), pH, and DO were continuously measured by logger probes. Total alkalinity (TA) and salinity were discrete measurements, with TA ranging from 2003.4 to 2043.3 μmol/kg and salinity ranging from 29.42 to 29.96 ppt throughout the course of the experiment. Refer to **Supplementary Table 1** for complete data table.

| treatment | temp (°C) | Target pH | pH | Target DO | DO (mg/L) |
| --- | --- | --- | --- | --- | --- |
| ambient pH ambient DO | 12.17 ± 0.17 | 7.85 | 7.83 ± 0.05 | 8.9 | 8.9 ± 0.2 |
| ambient pH low DO | 12.12 ± 0.12 | 7.85 | 7.85 ± 0.02 | 8.9 | 3.0 ± 0.1 |
| low pH ambient DO | 12.14 ± 0.08 | 7.45 | 7.45 ± 0.05 | 3.0 | 8.9 ± 0.1 |
| low pH low DO | 12.15 ± 0.16 | 7.45 | 7.49 ± 0.11 | 3.0 | 3.0 ± 0.3 |

### A combination of statistical methods reveal condition-specific responsive compounds

In general, individual metabolite and lipid profiles showed high variation regardless of treatment group, with no obvious trends revealed by clustering the relative abundance of all compounds by treatment groups (**Supplementary Fig. 2**). To identify specific compounds affected by treatments, we applied both univariate and multivariate statistical approaches, combining the strengths of different methods^27^. To assess individual effects as well as interaction effects from pH and DO treatments on individual compounds, we used a two-way analysis of variance (ANOVA) on the raw abundance data (**Supplementary Table 3 and 4**). Prior to performing ANOVA, we verified compounds showed homogeneity of variances across treatment groups (>96% compounds showed a Levene test *P* value > 0.05), and that metabolite and lipid abundances were mostly (on average 69% compounds showed a Shapiro-Wilks test *P* value > 0.05) normally distributed within treatment groups (**Supplementary Table 5**). Although about one-third of compounds violated the ANOVA normality assumption, ANOVA can be considered robust to violations of this assumption when datasets have more than 10 samples per treatment group^28,29^. Of all compounds analyzed, 56/651 metabolites (including 24/160 known metabolites) and 98/3095 lipids (including 7/284 known lipids) showed overall model significance at *P* < 0.1 and at least one model term (pH, DO, and/or pH x DO interaction) significant at *P* < 0.05 (**Supplementary Fig. 3 and 4, and Supplementary Table 6**). Because no metabolite or lipid overall model *P* values passed their respective 10% false discovery rate Benjamini-Hochberg *P* value threshold of 1 x 10^-4^ and less than 3 x 10^-5^, we chose to use their uncorrected *P* values.

To identify discriminatory compounds and model the relationship between metabolite profiles and exposure treatments using a multivariate linear regression approach, we used partial least square discriminatory analysis (PLS-DA). For facilitating comparisons of metabolite composition among treatment groups, compound abundances were centered around the mean and scaled by the reference group (ambient pH and DO) compound standard deviations prior to multivariate analysis^30^. The PLS-DA model for the metabolite data shows partial separation of low pH-treated samples from ambient pH-treated samples by the first and second components accounting for a total of 30.9% (**Fig. 1a**). Moreover, the metabolite data shows partial separation of low DO-treated samples in the second and third components, explaining an additional 6.8% of variation (**Fig. 1b**). For the lipid data, the PLS-DA model shows less clear treatment group separation on the first component (**Fig. 1c**) but shows partial separation of low pH-treated samples from ambient pH-treated samples on the second and third components explaining 14.7% of variation (**Fig. 1d**). Because the overall PLS-DA model had weak predictive power (**Supplementary Fig. 5**), we used importance thresholds defined by the point of diminishing returns in the PLS-DA component loadings plots (**Supplementary Fig 6.** and **Methods**), and identified a total of 45 metabolites (including 14/160 known metabolites) and 18 unknown lipids as compounds important in discriminating between treatment groups.

**Figure 1.**
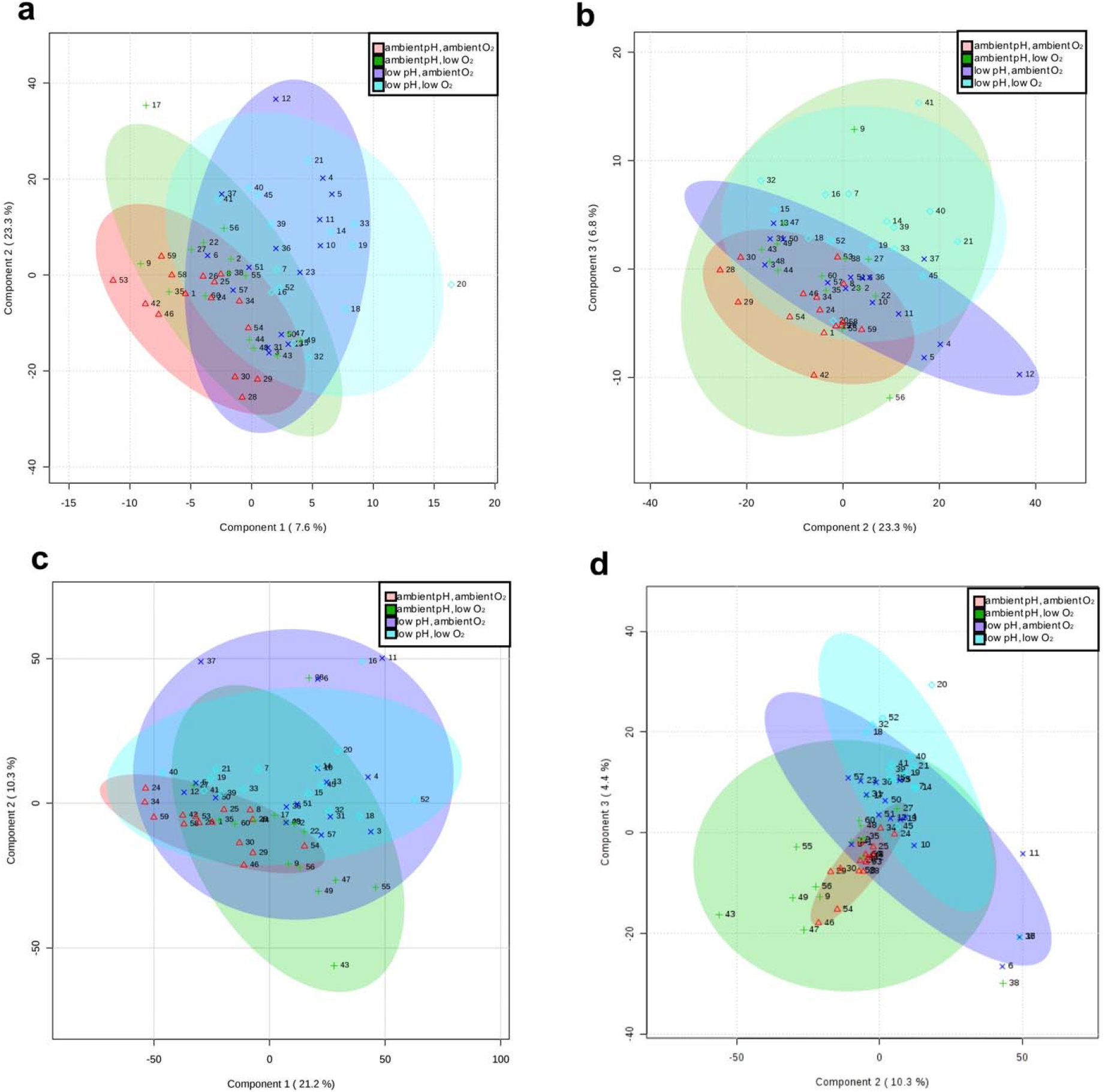
Summary of PLS-DA analysis. Metabolite data show partial separation of low pH treatment groups (blue and purple) from low DO treatment groups (pink and green) by (**a**) Components 1 and 2 and (**b**) components 2 and 3. Lipid data show partial separation of treatment groups by (**c**) components 1 and 2 and (**d**) components 2 and 3. Shapes correspond to treatment groups (ambient pH and DO, Δ; ambient pH and low DO, +; low pH and ambient DO, x; low pH and low DO, ◇. Numbers refer to the sample number.

Finally, we used a multivariate nonlinear-based supervised random forest classification to predict treatment classes from the metabolomic and lipidomic data (**Supplementary Fig. 7a-b**). A total of 9 metabolites (including 4/160 known metabolites) and 15 lipids (including 1/284 known lipids) were considered compounds important to predicting treatments based on how much their removal from the model decreased prediction accuracy (**Supplementary Fig. 7c-d, Supplementary Table 8**, and **Methods**). While no compounds were commonly identified by all 3 statistical methods as was expected given the fundamental differences of the methods, 7 metabolites were commonly identified by ANOVA and PLS-DA, 8 metabolites were in common between ANOVA and random forest, 1 metabolite overlapped between random forest and PLS-DA, and 4 lipids overlapped between ANOVA and random forest (**Fig. 2a-b**). This combination of analyses yielded a comprehensive set of compounds affected by treatments that no one method was capable of capturing on its own.

**Figure 2.**
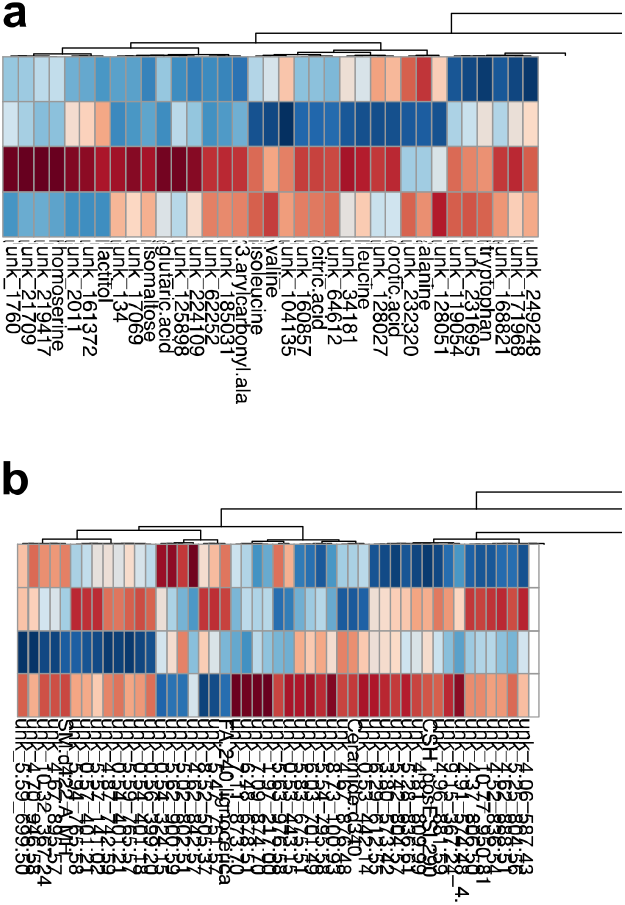
Venn diagrams DA, and/or random forest a

### Low oxygen more strongly affects compound abundance than pH

Heatmaps of the statistically identified 94 (including 35 known) metabolites and 127 (including 7 known) lipids, show several compounds respond to low pH and oxygen treatments by commonly increasing or decreasing abundance relative to the ambient treatment (**Fig. 3a-b**). This is also summarized by violin plots of known compound abundances in **Figure 4a-c**. For example, 5-methoxytryptamine, butyrolactam, cysteine, cysteine, homoserine, pipecolinic acid, piperidone, ribose, and xanthine commonly show a decrease in abundance in response to low DO treatments relative to ambient DO treatments. Glutamic acid, maltose, and maltotriose commonly show an increase in abundance in response to low DO treatments relative to ambient DO treatments (**Figure 4a**). Overall, metabolite abundance tends to decrease rather than increase in response to low DO treatments relative to ambient DO treatments, exemplified by the mostly blue color of compound abundance averages for the low DO treatment group (green) in **Figure 3a**. Specific to the low pH treatment, more metabolites show average abundances similar to ambient treatment signifying pH has a less dramatic effect on metabolites compared to low DO treatment (**Fig. 3a-b and Fig. 4a-b**). This is also apparent in the combined low pH and low DO treatment, where metabolites and lipids show average abundances similar to the low DO treatment signifying low DO has a more dominant effect on compounds than low pH (**Fig. 3a-b**).

**Figure 3.**
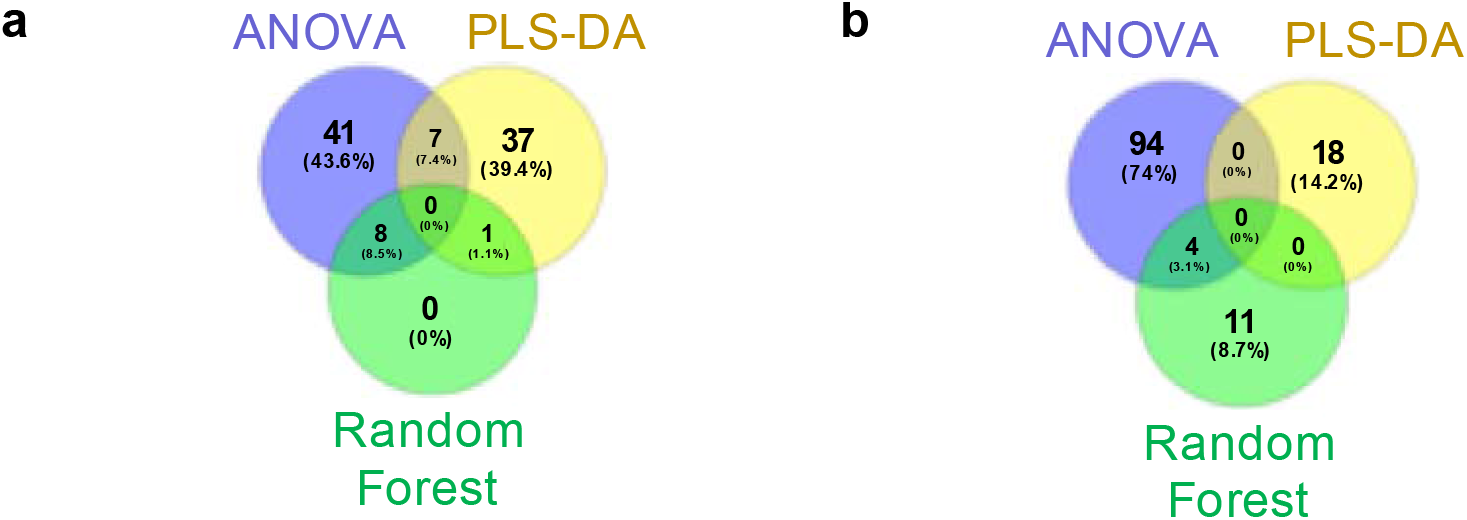
Heatmap plots showing the average compound abundance (average peak intensity) for each treatment group for the (**a**) 94 general metabolites and (**b**) 127 lipids selected by multivariate and univariate statistical methods used to evaluate treatment effects. Individual known and unknown (“unk”) compound names are listed over the columns and treatment groups are listed over the rows (orange, ambient pH, ambient O_2_; green, ambient pH, low O_2_; purple, low pH, ambient O_2_; and blue, low pH, low O_2_). Compound average abundances are shown as auto-scaled within each compound (“relative abundance”).

**Figure 4.**
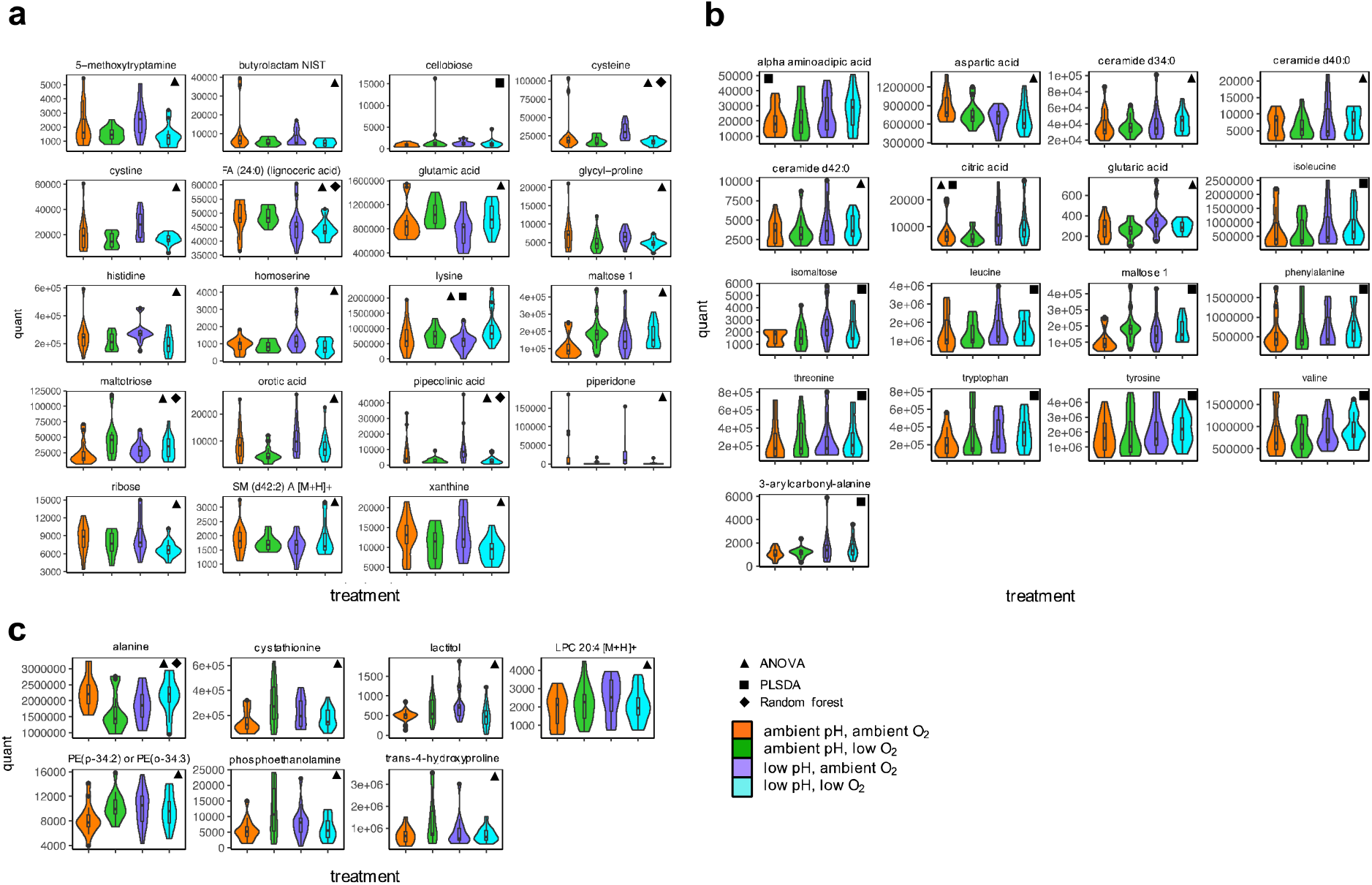
Compound abundance levels (peak intensities) of known metabolites and lipids selected by univariate and multivariate statistical methods used to evaluate treatment effects. Violin plots with boxplot insets show the distribution of compound abundances within each treatment group for compounds statistically showing (**a**) DO treatment effect, (**b**) pH treatment effect, and (c) DO:pH interaction effect. Binbase names for each known compound are listed across the top of each plot. Abundance levels (peak intensities, noted as “quant”) are listed on the y-axis while treatments are listed along the x-axis. Statistical methods that each known compound was identified by are noted by shapes in the upper corners of each plot (ANOVA, triangle; PLS-DA, square; and random forest, diamond).

### Pathway analysis reveals potential mechanisms of pH and oxygen stress tolerance

To explore the physiological relevance of treatment-responsive compounds, we performed biochemical pathway analysis on all known compounds. While compounds classified as sugars and fatty acids were affected by the low pH and DO treatments, most of the significantly affected compounds were part of the amino acids class, indicating amino acid metabolism was most significantly altered by the treatments. Abbreviated biochemical pathway networks focusing on affected amino acids are shown in **Figure 5**, and the complete biochemical pathway networks affected by low pH, low DO, and the combined low pH and DO treatments can be found in **Supplementary Figures 8-10**. The trends in amino acid abundances resulting from either low pH, low DO, or the combined low pH and DO treatment (**Fig. 5a-c**), suggest different amino acid metabolic pathways are affected in response to each factor or combination of factors.

**Figure 5.**
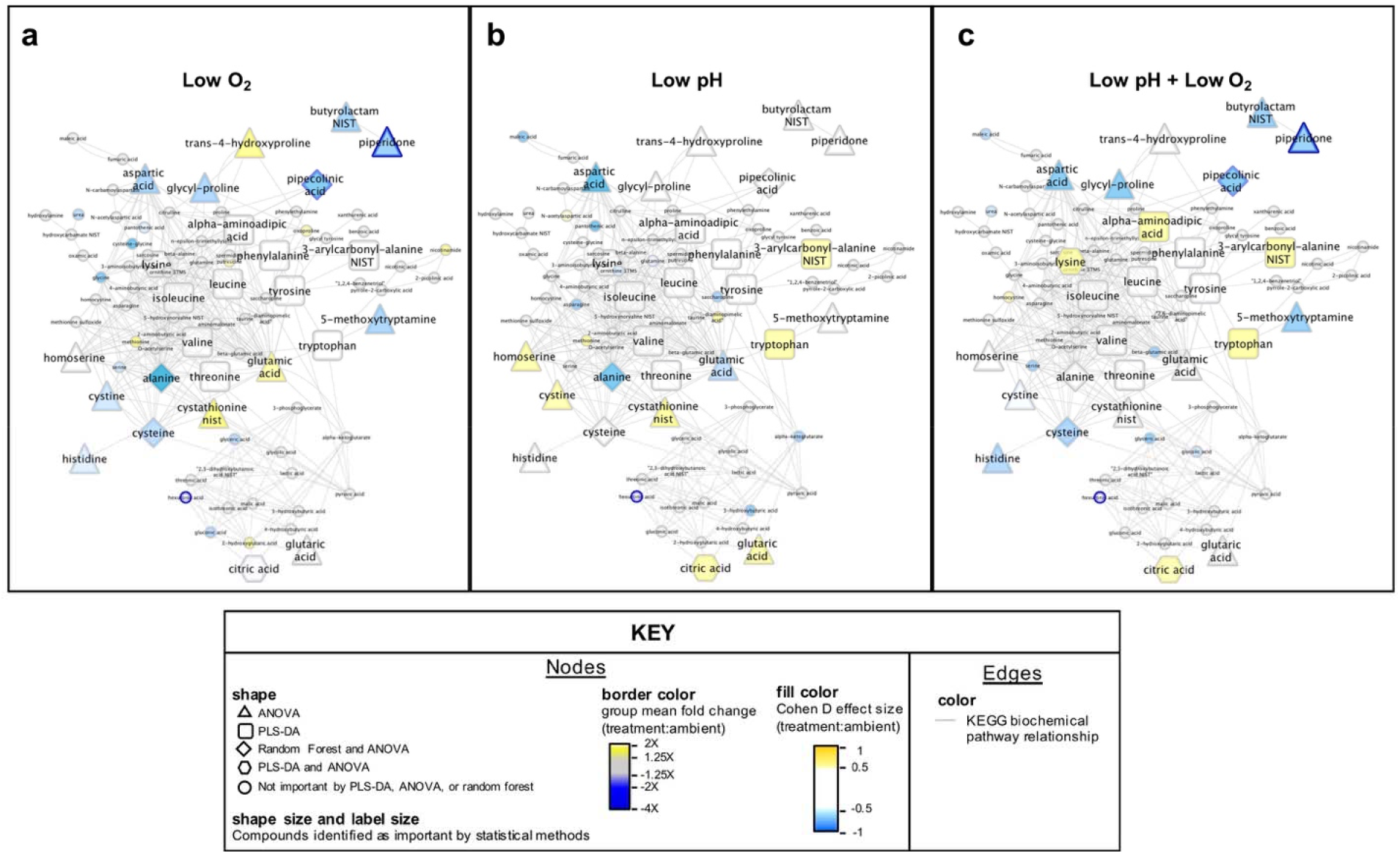
Differential amino acid network effects from (**a**) low DO, (**b**) low pH, and (**c**) combined low pH and low DO. Compounds are clustered by chemical similarity. Node fill color is colored by Cohen D effect size comparing the treatment group to the ambient pH,ambient DO group, and node borders are colored by the treatment group mean fold change relative to the ambient pH,ambient DO group. Node shapes indicate the statistical method(s) from which the compound was classified as important. Node shape and label size are enlarged if the compound was identified as important by a statistical method. Gray edges indicate nodes sharing a KEGG biochemical pathway(s).

Specific to low DO treatment response compared to the normoxic treatment, energy conservation pathways appear to be most affected (**Fig. 5a**). The increased lysine with decreased pipecolinic acid and the piperidine derivative piperidone abundance is suggestive of downregulation of the lysine degradation pathway^31,32^. The decreased abundance of both cysteine and the cysteine homodimer, cystine, suggests that cysteine catabolism is upregulated. This coupled with the increased abundance of glutamic acid suggests the glutathione synthesis pathway could be downregulated, as both cysteine and glutamic acid are precursor molecules in glutathione synthesis^33,34^. Glutathione itself was not detected, but is typically challenging to detect by MS techniques in marine animals due to its reactivity^35^. The glycogen intermediates maltose and maltotriose show an increase in abundance consistent with previously observed glycogen synthesis pathway upregulation during hypoxic stress^36^. Purine and pyrimidine metabolic intermediates orotic acid, ribose, and xanthine show decreased abundance suggesting purine and pyrimidine metabolic pathways could be downregulated^37^.

Specific to low pH treatment response compared to ambient pH treatment, ATP generation pathways appeared most affected (**Fig 5b**). The decreased abundance of aspartic acid and the subtle decreased abundance of maleic acid may suggest the citric acid cycle intermediates that they form (oxaloacetate, malic acid, and fumaric acid)^38,39^ are favored. Although, oxaloacetate was not detected (likely due to the instability of the alpha keto acid compound)^40^, and malic acid and fumaric acid did not show a significant difference across treatments. However, citric acid did show an increase in abundance. Taken all together, the altered abundance of these compounds suggest that citric acid cycle activity could be upregulated in response to low pH^41^. An increase in glutaric acid in response to low pH suggests the catabolism of its parent molecule, glutaryl CoA, which produces ATP in addition to glutaric acid^42^. Glutaryl CoA catabolism also supports citric acid cycle activity because glutaryl CoA can uncompetitively inhibit the alpha-ketoglutarate dehydrogenase complex that facilitates a rate limiting step in the citric acid cycle^43^.

Among the compounds affected by the combined low pH and DO treatment, phosphoethanolamine and phophatidylethanolamine (PE(p-34:2) or PE(o-34:3)) show increased abundance compared to ambient treatment (**Supplementary Fig. 10**), suggesting that low pH and DO might alter the glycerophospholipid synthesis pathway of which phosphatidulethanolamine is a product and phosphoethanolamine is a substrate intermediate in a rate limiting step^44^. Alanine shows a decrease in abundance, suggesting that alanine synthesis is decreased and that its substrate for synthesis, pyruvate, may be limited^45^. The increased abundance of cystathionine and subtle increase of methionine suggest cysteine and homoserine synthesis are likely stalled^46^ in response to combined low pH and low DO exposure. This overlaps with the decrease in homoserine abundance and decrease in cysteine synthesis products (cysteine and cystine) observed in low DO treatment (**Fig. 5b-c**).

## DISCUSSION

We used metabolomics to explore how the Dungeness crab might respond to the simultaneous change in oxygen and carbon dioxide in predicted future climate change scenarios in the Pacific Northwest. Although the untargeted metabolomics and lipidomics approaches rapidly identified hundreds of different molecular species, the majority of compounds identified in our study, including most compounds selected by statistical methods, lack annotations in existing metabolite databases. Thus, our understanding of their role in response to low pH and low DO treatment is limited without extensive further validation. Still, through shedding light on the simultaneous activity of hundreds of known compounds, we were able to observe a dynamic range of metabolic responses among individuals within treatment groups that indicates that Dungeness crab have flexibility in how their biochemistry compensates for environmental change. While we attempted to control for variation between individuals by collecting animals from the same location within a 2-month period, we were unable to control for prior environmental exposure or genetic background of the wild-caught animals we used. Where prior studies have found high genetic variation among individuals within one sampling site^47,48^, we suspect that both genetics and prior environmental exposure likely contributed to the high variation in metabolite abundances among individuals within treatment groups, which may have obscured the different treatment effects. Within-sample variation could be reduced by sampling from a specific tissue rather than using whole animals, however this may be challenging in early stage juvenile Dungeness crabs given their small size.

We applied three different statistical methods (ANOVA, PLS-DA, and random forest) for selecting important metabolites in order to combine the strengths of powerful univariate and multivariate analyses^27^. Although using univariate statistics in the strictest sense with an FDR correction to control for false discoveries showed no statistically significant variation for compounds across treatments, not all compounds were identified with the same confidence level and were subject to the sensitivity of the mass spectrometry method used. Applying an FDR correction to all detected compounds assumes that all compounds had equal chance for discovery, which can mask important biology in highly variable datasets^49^. For this reason and due to our use of wild-caught whole-animal samples, we chose a more liberal focus on all compounds showing an overall ANOVA model *P* value < 0.1 and at least one model term *P* value of < 0.05, or identified as important in PLS-DA or random forest models in our pathway analysis.

While we acknowledge a level of uncertainty in our pathway analysis due to liberal selection of important treatment-responsive compounds, the resulting proposed affected pathways are consistent with previous observations of pH and hypoxia effects on different organisms. In general, amino acid metabolism is a well-documented mechanism for stress tolerance^50,51^. This class of compounds contains versatile chemical structures that serve as buffering molecules, antioxidants, signaling molecules, and chemical building blocks for the synthesis of proteins important in stress response (i.e., heat shock proteins, unfolded protein response proteins, ion channels). Under low oxygen conditions, it is in the best interest of the animal to limit non-essential energy consuming pathways^52^. One way Dungeness crab may do this is through reducing the activity of evolutionarily conserved gamma-glutamyl cycle that synthesizes glutathione and consumes ATP^53,54^ by limiting cysteine availability through catabolism. Interestingly, glutathione reduction in response to hypoxia has been observed in multiple mammalian cell lines^55–58^. Also under low oxygen conditions in nature, food can be scarce, and it is theorized that glycogen storage during low oxygen can help prepare cells for low nutrient conditions. In multiple mouse and human cancer cell lines^36,59,60^, increased glycogen storage has been observed in response to hypoxia, and is induced by highly evolutionarily conserved hypoxia-inducible factor transcriptional signalling^36^. It seems that Dungeness crab may also adopt this strategy in combating low oxygen conditions. Under low pH, adult Dungeness crab initially develop hypercapnia which then abates over time via elevated hemolymph bicarbonate likely generated through gill restructuring and the upregulation of energy consuming ion-exchange proteins^12^. Our results indicate that this process may occur in early juveniles in low pH conditions given that we found metabolite profiles that support energy generation via increased citric acid cycle activity.

Having parsed out effects from low oxygen and low pH, we found that in combined low oxygen and pH conditions low oxygen has a more dramatic effect on metabolite abundance. However, the animals in this treatment group still showed pathway alterations similar to the low pH treatment group including increased abundance of citric-acid-cycle-related metabolites to potentially increase ATP production. It is not yet clear what the longer-term consequences are of these metabolic adjustments or how long these responses could be sustained. Future avenues of research to expand on these findings should include targeted metabolomics to confirm the compounds identified in this study as well as capture more antioxidants and citric acid cycle intermediates that were not detected in this study. Targeted expression profiling would also be helpful in confirming the biochemical pathway activity predictions from this study. Ultimately, longer-term exposure experiments on crabs reared over generations might best reveal the maximal duration that these metabolic responses can be sustained and any long-term consequences of low pH and oxygen exposure. This exploratory metabolomics and lipidomics analysis uncovered potential biochemical pathways affected by experiments simulating ocean acidification and hypoxia, and can now serve as preliminary hypotheses for deeper investigations of how the Dungeness crab (and even other crustaceans) may tolerate global ocean change.

## METHODS

### Animals

*Cancer magister* megaolopae were collected from a single site in Puget Sound (47.950232, - 122.301784) on several days between the June and September 2016 using light traps that were set overnight. The contents of each trap were immediately transferred into a 5-gallon bucket that was, within 5 minutes, pooled into a cooler with ice packs and an air bubbler. After transferring contents from 7 traps in approximately 1 hour, the cooler containing megalopae was brought within 5 minutes into the lab and megalopae were individually transferred onto the water flow-through system described below. The total time for transferring megalopae from the cooler onto the seawater flow-through system was about 2 hours.

### CO_2_ exposure experiments

Megalopae were held in individual 250 mL customized jars on Mobile Ocean Acidification Treatment Systems (MOATS). These systems flowed one-micron-filtered, UV-sterilized, Puget Sound seawater maintained at 12°C. Prior to flowing through jars, seawater was degassed and oxygen, nitrogen and carbon dioxide gases were resupplied to finely control dissolved gas levels. Temperature, pH, and dissolved oxygen were continuously monitored throughout the duration of the experiment by Omega thermistors, Honeywell Durafet III probes, and Vernier optical dissolved oxygen probes, respectively. The pH was additionally validated by periodic sampling of water for dissolved inorganic carbon and total alkalinity, and by bi-weekly spectrophotometric pH measurements using an Ocean Optics USB 230 2000+ Fiber Optic Spectometer with SpectraSuite software and a 5mM solution of Sigma Aldritch m-cresol purple indicator dye. MOATS chemistry parameters were automatically adjusted through a data-driven feedback system. Megalopae were fed *Artermia salina* (San Francisco Bay brand) at a target concentration of 1 nauplius per mL every 3 days. Once megalopae transitioned to the first juvenile instar, they were fed small pieces of squid. Upon transitioning to the second juvenile instar, crabs were held on the MOATS for an additional 48 hours post-molt in attempt to reduce variation due to potential stochastic physiological processes associated with molting ^61,62^. Juveniles were then immediately frozen and stored at −80°C after lightly blotting with a paper towel to remove excess seawater. To reduce variation from length of exposure, which in total ranged from 20-65 days, only 60 crabs with the average exposure time (30-33 days) were chosen for metabolomics analysis with 15 individuals from each treatment group.

### Water chemistry analysis

Mean and standard deviation calculations for pH, DO, and temperature (**Table 1**) were based on logger data (measured every 10 minutes) with the exception of pH and DO data from MOATS 5 and 12, where Spectrophotometric pH and Presens DO measurements were instead used due to incorrectly calibrated pH and DO probes in these MOATS. Temperature logger readings above 25°C and below 4°C were excluded from the mean and standard deviation analysis because these were out of the achievable range for the equipment used and were indicative of a thermistor malfunction. Total alkalinity (TA) and salinity were discrete measurements, with TA ranging from 2003.4 to 2036.1 μmol/kg and salinity ranging from 29.42 to 29.96 ppt throughout the course of the experiment. The complete data table is included as **Supplementary Table 1**.

### Sample preparation

Frozen samples were sent to the West Coast Metabolomics Center, Davis, CA for sample preparation, metabolomics and lipidomics profiling. Whole animals were thawed, weighed, and derivatized as previously described^63,64^ (**Supplementary Table 2**). Briefly, samples were extracted at −20°C with 2 mL of degassed acetonitrile/isopropanol/water (3:3:2) solution and solvents were evaporated to complete dryness with a Labconco Centrivap cold trap concentrator. Membrane lipids and triglycerides were subsequently removed from dried samples with 50% acetonitrile, and samples were again concentrated to complete dryness. 15 mg of each sample preparation was used for metabolomic profiling. For lipidomic profiling, 15 mg of the same sample preparation was also used to which internal standards, C8-C30 fatty acid methyl esters were added. Aliquoted samples were derivatized with methoxyamine hydrochloride (Sigma-Aldrich) in pyridine (Acros Organics) and then by N-methyl-N-(trimethylsilyl) trifluoroacetamide (Sigma-Aldrich) for trimethylsilylation of acidic protons.

### Metabolite and lipid data acquisition

General metabolite and lipid abundances were quantified from derivatized samples by gas-chromatography, time-of-flight mass spectrometry (GC-TOF/MS) and charged-surface, hybrid-column, electrospray-quadrupole, time-of-flight mass spectrometry (CSH-ESI QTOF MS/MS), respectively. For metabolites, an Agilent 6890 gas chromatograph (Santa Clara, CA) was used with a Leco Pagasus IV time-of-flight mass spectrometer running Leco ChromaTOF software 2.32 (St. Joseph, MI). The following temperature profile was used: 50°C to 275°C final temperature at a rate of 12°C/s and hold for 3 minutes. Injection volume was 0.5 μl with 10 μl/s injection speed on a splitless injector with a purge time of 25 seconds. Liner (Gerstel #011711-010-00) was changed after every 10 samples, (using the Maestro1 Gerstel software vs. 1.1.4.18). Before and after each injection, the 10 μL injection syringe was washed 3 times with 10 μL ethyl acetate. For gas chromatography, a 30 m long, 0.25 mm i.d. Rtx-5Sil MS column (0.25 μm 95% dimethyl 5% diphenyl polysiloxane film) with additional 10 m integrated guard column was used (Restek, Bellefonte PA). 99.9999% pure Helium with a built-in purifier (Airgas, Radnor PA) was set at a constant flow of 1 mL/minute. The oven temperature was held constant at 50°C for 1 minute and then ramped at 20°C/minute to 330°C at which it is held constant for 5 minutes. The transfer line temperature between gas chromatograph and mass spectrometer was set to 280°C. Electron-impact ionization at 70V was employed with an ion source temperature of 250°C. Acquisition rate was 17 spectra/second, with a scan mass range of 85-500 Da.

For positively charged lipids, an Agilent 6530 QTOF mass spectrometer with resolution 10,000 was used and for negatively charged lipids, an Agilent 6550 QTOF mass spectrometer with resolution 20,000 was used. Electrospray ionization was used to ionize column elutants in both positive and negative modes. Compounds were separated using a Waters Acquity ultra-high-pressure, liquid-chromatography charged surface hybrid column C18 (100 mm length x 2.1 mm internal diameter; 1.7 um particles) using the following conditions: mobile phase A (60:40 acetonitrile:water + 10 mM ammonium formiate + 0.1% formic acid, mobile phase B (90:10 isopropanol:acetonitrile + 10 mM ammonium formiate + 0.1% formic acid), 65°C column temperature, a flow rate of 0.6 mL/minute, an injection volume of 3 uL, an injection temperature of 4 C, and a gradient of 0 minutes 15%, 0-2 minutes 30%, 2-2.5 minutes 48%, 2.5-11 minutes 82%, 11-11.5 minutes 99%, 11.5-12 minutes 99%, 12-12.1 minutes 15%, and 12.1-15 minutes 15%. The capillary voltage was set to +3.5 and −3.5 kV, and the collision energy to 25 or 40 eV for positive and negative modes. Mass-to-charge ratios (m/z) were scanned from 60 to 1700 Da and spectra acquired every 2 seconds.

### Spectral data processing

Acquired metabolite data were processed using UC Davis’s BinBase workflow, which performs data processing including peak detection at signal-to-noise levels of 5:1 throughout the chromatogram. Resulting apex masses are reported for use in the BinBase algorithm to facilitate metabolite identification and quantification^65^. Metabolites were identified through comparison to the BinBase database^22^ and peak heights were normalized to total metabolite content^66^.

For acquired lipid data, MassHunter (Qual v. B05.00) was used to find peaks from the raw data in up to 300 chromatograms. These peaks were imported into Mass Profiler Professional for alignment to determine which peaks occur in at least 30 % of the chromatograms. These peaks were then quantified with MassHunter. Resulting accurate mass data and tandem MS/MS spectra were compared to LipidBlast libraries for compound identification^21^. All spectra mapping to previously identified compounds are accessible in the public database MassBank of North America (http://mona.fiehnlab.ucdavis.edu/). Spectra that did not map to previously identified compounds are accessible by searching their BinBase identifiers listed in **Supplementary Tables 3 and 4** in the BinVestigate database (http://binvestigate.fiehnlab.ucdavis.edu).

### Statistical analyses

All statistical analyses excluded 18 compounds that were detected in less than half of the individual crabs surveyed, and were carried out in R, except for PLS-DA and random forest analyses which were carried out using the MetaboAnalyst web interface.

### Univariate statistics

Prior to applying a univariate statistical test, normality and heteroscedasticity of compound abundances within treatment groups was first assessed using the Shapiro-Wilks test and the Levene test, respectively (**Supplementary Table 5**). After confirming normality and heteroscedasticity assumptions were correct, a two-way ANOVA was then applied to each compound. A Benjamini-Hochberg FDR correction was applied to ANOVA *P* values to correct for multiple testing, but corrected *P* values were not used in selecting important compounds. Metabolites and lipids were selected for pathway analyses if they had an overall model ANOVA *P* value less than 0.1 and an effect *P* value with less than 0.05 without an FDR correction applied. Results from ANOVA tests are reported in **Supplementary Table 6**.

### Multivariate statistics

Prior to applying multivariate testing, data were normalized by mean-centering and scaling by the reference-group standard-deviations of each compound in order to make compounds more comparable to one another^30^. Normalized data was then uploaded to MetaboAnalyst with no further normalizations applied. The PLS-DA function was run with default settings and a table of component loadings for each compound were exported from MetaboAnalyst (**Supplementary Table 7**). Compound loadings were ordered by from greatest variance explained to lowest, and plotted for PLS-DA components. Importance thresholds were placed at the point of diminishing returns in the plot curves (**Supplementary Fig. 6**). Specifically for metabolites, loadings were plotted PLS-DA components 1, 2, and 3 (**Supplementary Fig. 6a-c**) since these PLS-DA components gave the greatest separation between treatment groups (**Fig. 1a-b**). The points of diminishing returns that importance thresholds were placed in the metabolite loadings plots were as follows: component 1, loadings threshold > 0.05; component 2, loadings threshold > 0.1; and component 3, loadings threshold > 0.05. Specifically for lipids, loadings were plotted for PLS-DA components 2 and 3 (**Supplementary Fig. 6d-e**) since these gave the best separation between treatment groups, while component 1 could only distinguish the ambient from the altered treatment groups (**Fig. 1c-d**). The points of diminishing returns that importance thresholds were placed in the lipids loadings plots were as follows: component 2, loadings threshold > 0.05; and component 3, loadings threshold > 0.05. For the random forest analysis, the Random Forest function was run with 2000 decision trees and either 25 predictors for the metabolite dataset or 56 predictors for the lipid dataset, conforming to the default classification value being the square root of the number of variables^67^. The complete table of important variables and their mean decrease in random forest accuracy prediction exported from MetaboAnalyst (**Supplementary Table 8**). The mean decrease in prediction accuracy for each compound was ordered from largest to smallest and plotted (**Supplementary Fig. 7c-d**). Importance thresholds were drawn from the plots at the points of diminishing returns, which for metabolites was a mean decrease in accuracy > 0.001 and for lipids was a mean decrease in accuracy > 0.00047.

Heatmaps in **Figure 3** and **Supplementary Figure 2** were made using the heatmap function in the MetaboAnalyst web interface with Euclidean distance, ward clustering, plotting auto-scaled compound abundance averages for each treatment group.

### Pathway analysis

Pubchem and KEGG identifiers were obtained for metabolites and lipid international chemical identifier keys provided by the West Coast Metabolomics Core using Chemical Translation Service^68^ and Pubchem Identifier exchange^69^. Metamapp^70^ was then used to map metabolites by chemical and biochemical relationships. The generated SIF file (**Supplementary File 1**) was imported into Cytoscape^71^. For visualizing the pH, DO, and pH:DO interaction effects, both mean abundance fold change and Cohen D effect size values were calculated for each compound within a treatment group relative to the ambient treatment group. Cohen D effect size 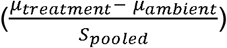 was calculated for each compound raw abundance within a treatment group relative to the ambient treatment group using the R package effsize^72^. The treatment group -log_2_ fold change values and the Cohen D effsize output from R were added to a node attribute file (**Supplementary Table 9**) that was also imported into Cytoscape. Low pH, low DO, and low pH:low DO networks were styled using the following settings: node shapes were set according to statistical method used to select the compound (ANOVA, triangle; PLS-DA, square; random forest, diamond; both PLS-DA and ANOVA, hexagon; not selected as important by any method, circle); node border was color by the treatment group mean compound abundance fold change relative to ambient (1.25X-2X fold change, yellow gradient’ −1.25X – 1.25X fold change, grey; −1.25X - -10X fold change, blue gradient); node fill was colored by the Cohen D effect size of treatment on compound abundance relative to ambient (effect size of 0.5-1, yellow gradient; −0.5.-0.5, white; and −0.5- −1, blue gradient); node shape size and label size were set to fixed values (150 height and width with size 50 font for statistically selected compounds, and 30 height and width with size 12 font for compounds not identified by statistics (**Supplementary Fig. 8-10**).

## DATA AVAILABILITY

All mass spectra mapping to previously identified compounds are accessible in the public database MassBank of North America (http://mona.fiehnlab.ucdavis.edu/). Spectra that did not map to previously identified compounds are accessible by searching their BinBase identifiers in the BinVestigate database (http://binvestigate.fiehnlab.ucdavis.edu). Processed metabolite and lipid spectral data are included as **Supplementary Table 3 and 4**.

## ACKNOWLEDGEMENTS

This material is based upon work supported by the Ocean Acidification Program and Northwest Fisheries Science Center of the National Oceanic and Atmospheric Administration, and in part by the National Science Foundation Graduate Research Fellowship Program and Graduate Research Internship Program (to S.A.T.). S.A.T. is supported in part by the Mary K. Chapman Foundation. We thank the current and former members of the Ocean Acidification lab at the Northwest Fisheries Science Center Mukilteo Research Station, members of the West Coast Metabolomics Core and Fiehn lab at University of California Davis.

## AUTHOR CONTRIBUTIONS

K.M.N., P.M., and D.S.B conceived the project. K.M.N., P.M., and D.S.B. advised research. M.M. and D.P. performed experiments. S.A.T. performed statistical analyses and pathway analysis. S.A.T. prepared the manuscript with edits from K.M.N., P.M, and D.S.B.

