## Supplementary Figures for "Uncovering mechanisms of global ocean change effects on the Dungeness crab (*Cancer magister*) through metabolomics analysis"

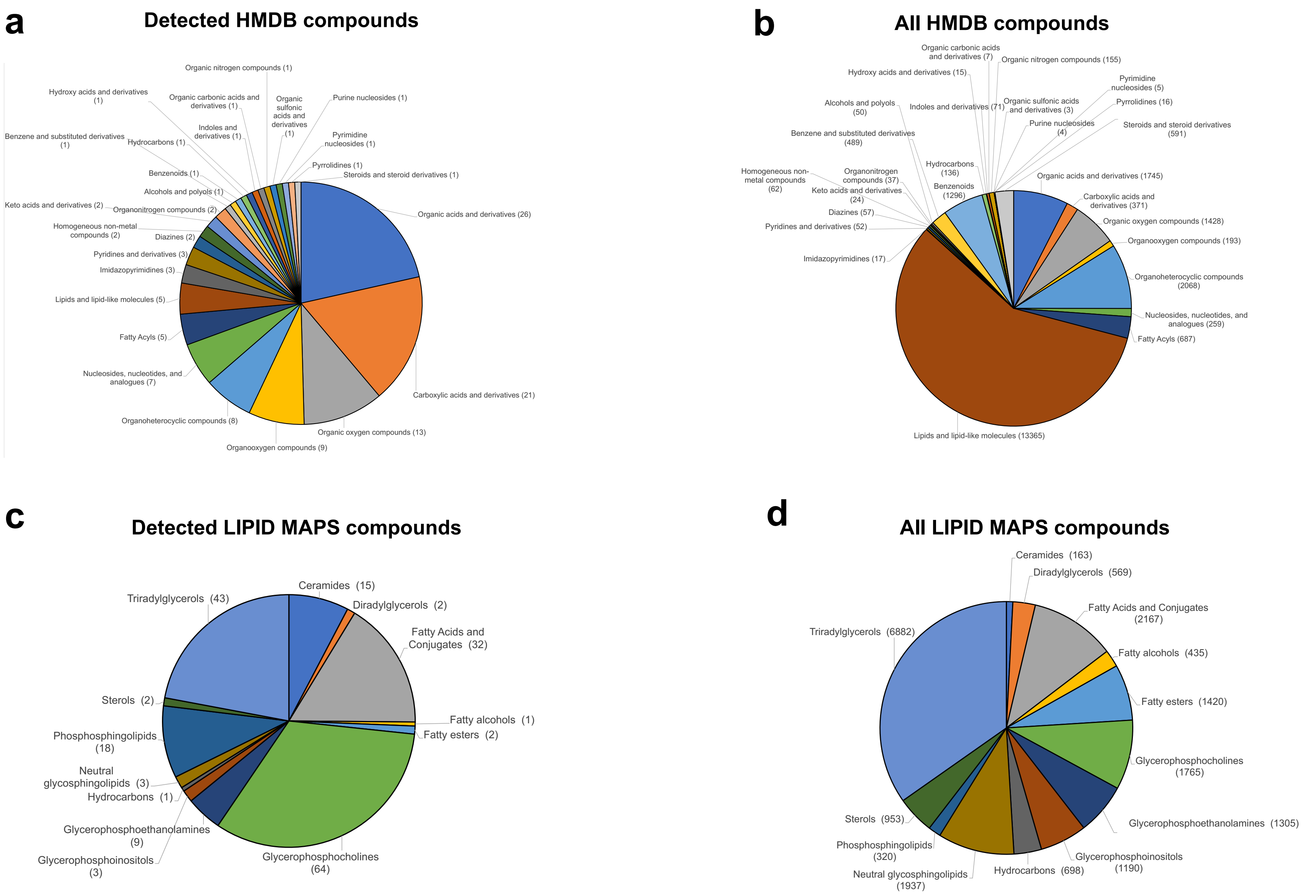


**Supplementary Figure 1.** Comparison of the distribution of compound classes for metabolites and lipids identified by mass spectrometry methods from juvenile Dungeness crab samples to the distribution of all compound classes listed in HMDB and LIPID MAPS databases. (**a**) Detected compounds that match entries with class information in the Human Metabolome Database (HMDB). (**b**) All compounds with class information listed in HMDB. (**c**) Detected compounds that match entries with class information in the LIPID MAPS database. (**d**) All compounds with class information listed in LIPID MAPS database.

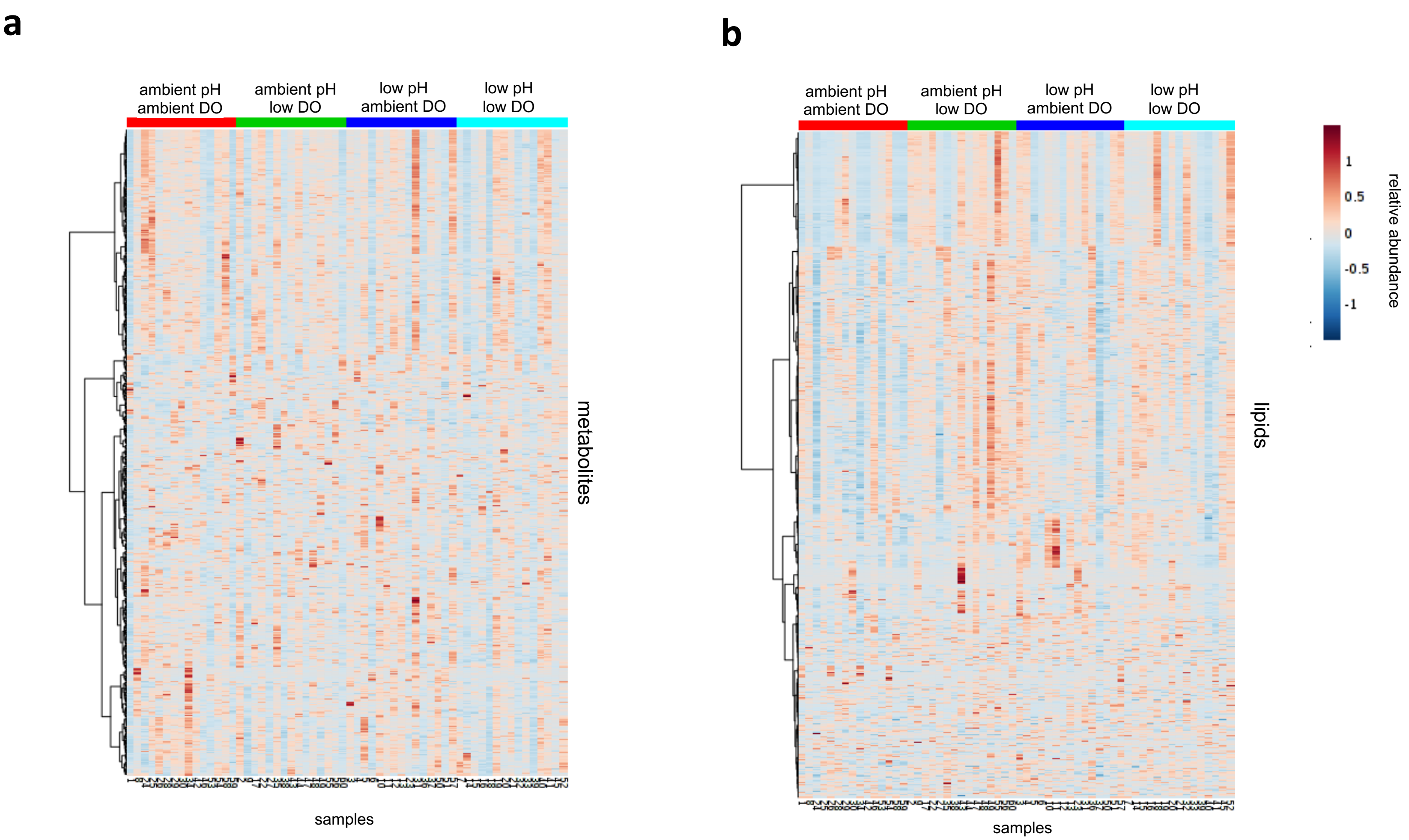


**Supplementary Figure 2.** Heatmap plots of (**a**) metabolite and (**b**) lipid relative abundances clustered by treatment group. Samples and treatment groups are listed along the x-axis and individual compounds are listed along the y-axis. Relative compound abundances are calculated by centering around mean compound abundance and auto-scaling. Dark red indicates high abundance and dark blue indicates low abundance.

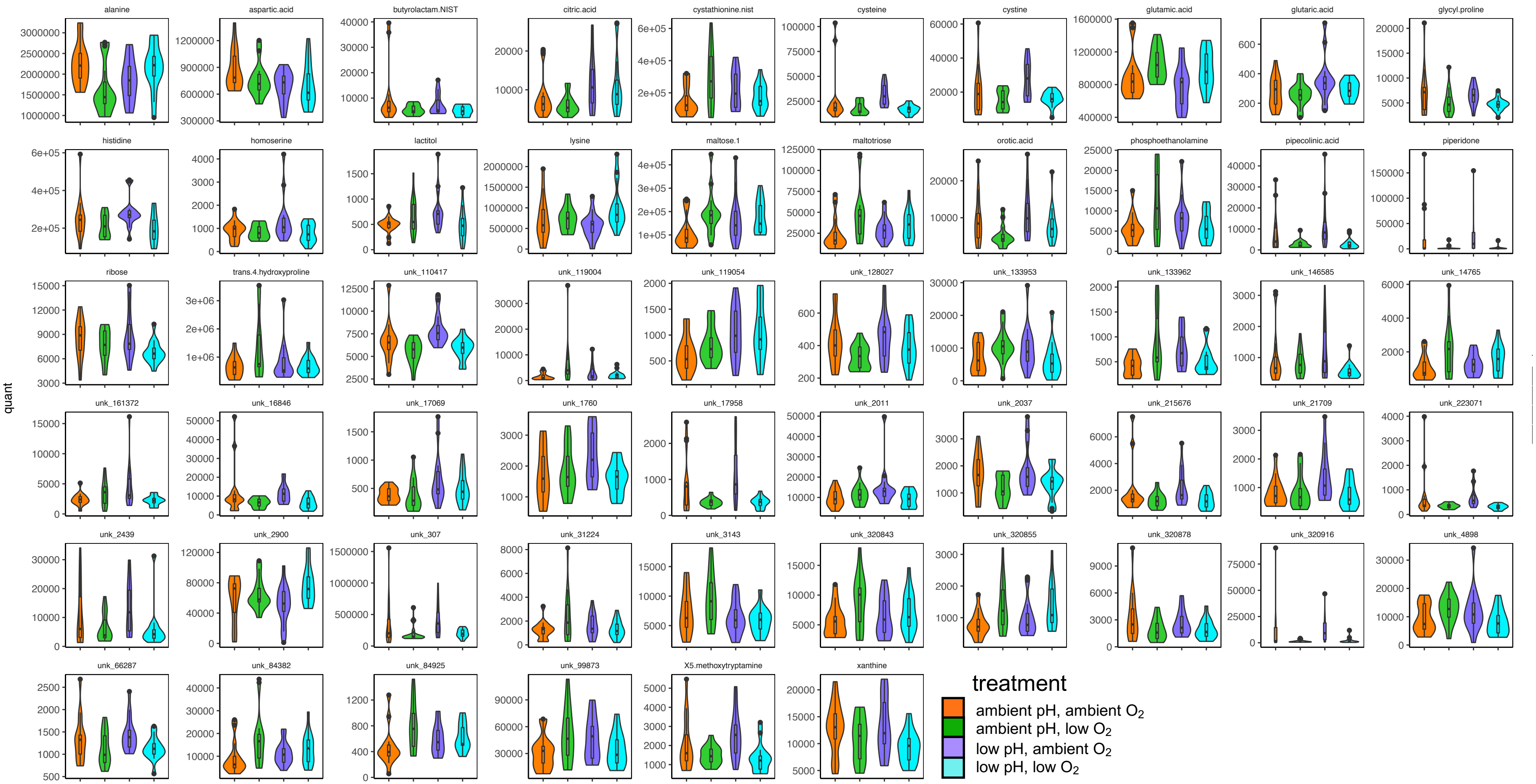


**Supplementary Figure 3.** Violin plots with boxplot inlays of metabolite abundances for metabolites with an overall ANOVA model *P* value < 0.1 and at least one ANOVA model term *P* value of < 0.05.

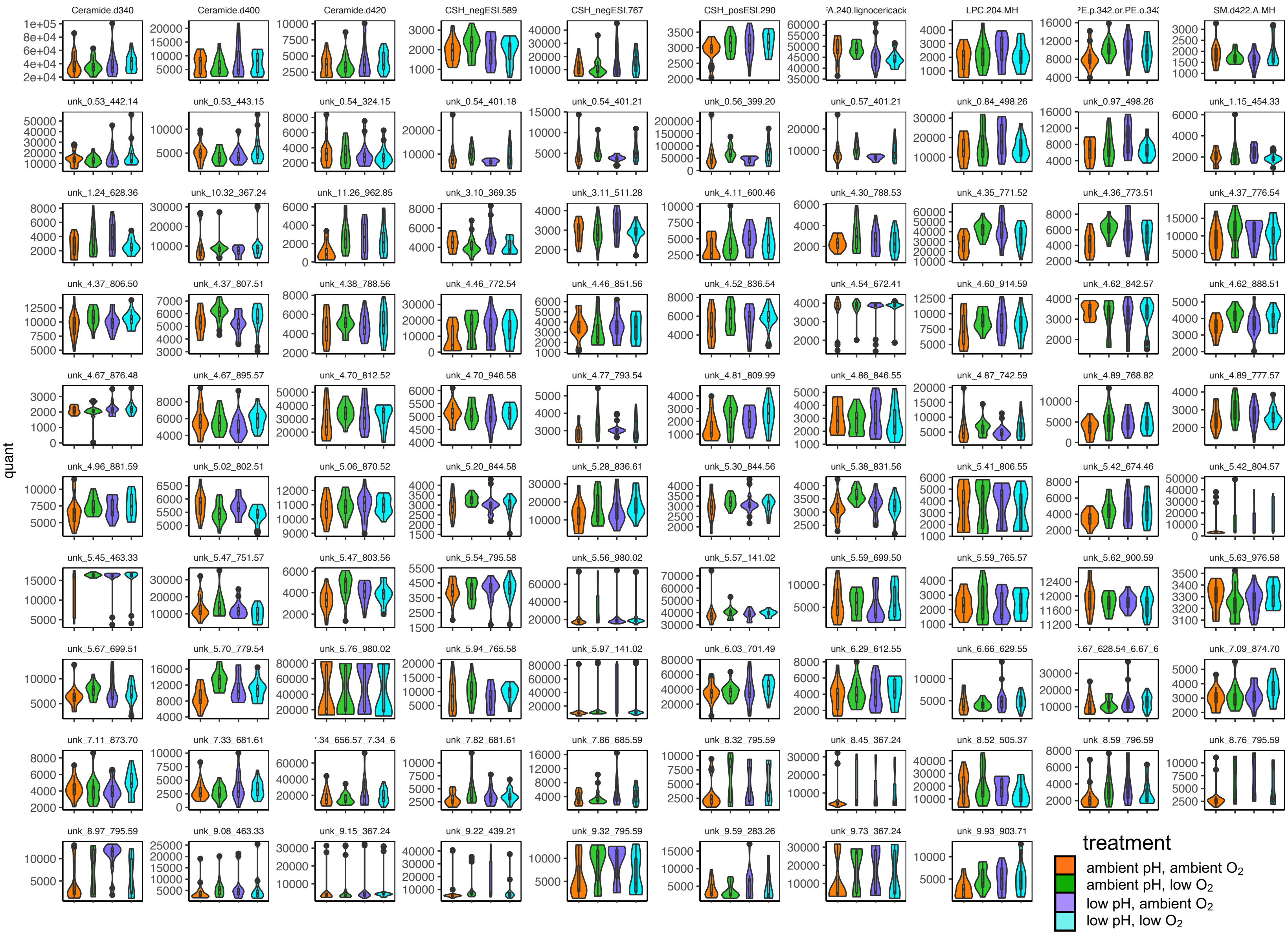


**Supplementary Figure 4.** Violin plots with boxplot inlays of lipid abundances for lipids with an overall ANOVA model *P* value < 0.1 and at least one ANOVA model term *P* value of < 0.05.

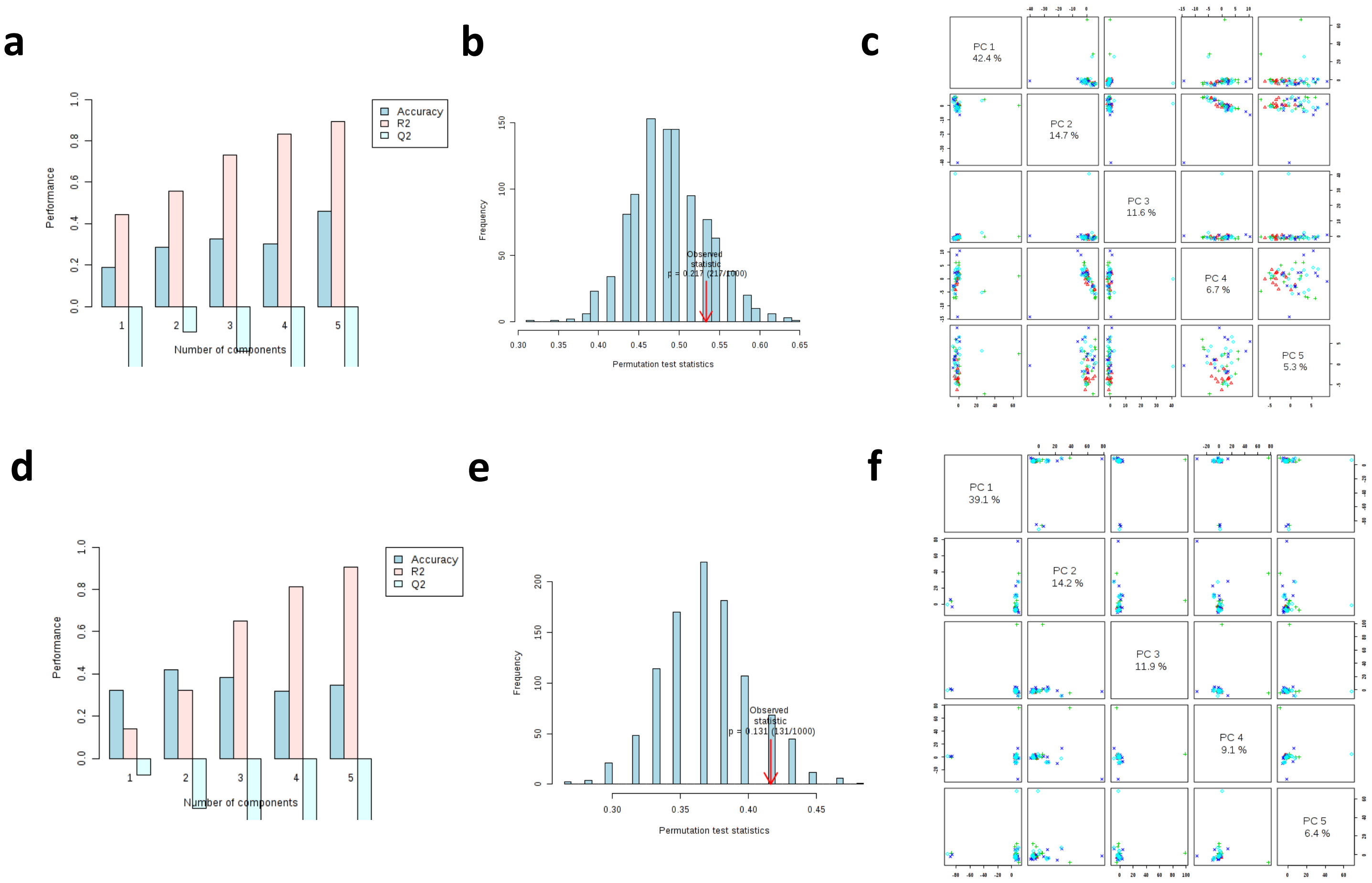


**Supplementary Figure 5**. PLS-DA validation. **(a)** QC accuracy plot for metabolite dataset. (**b**) Metabolite PLS-DA model performance plot compared to a PLS-DA model generated by 1000 permutations of randomly reassigned group labels. (**c**) Plots of the first 5 principal components of a PCA on the metabolites selected by PLS-DA. Colored shapes indicate treatment groups as follows: red triangle, ambient pH:ambient DO; green cross, ambient pH:low DO; blue X, low pH:ambient DO; turquoise diamond, low pH:low DO. **(d)** QC accuracy plot for lipid dataset. (**e**) Lipid PLS-DA model performance plot compared to a PLS-DA model generated by 1000 permutations of randomly reassigned group labels. (**f**) Plots of the first 5 principal components of a PCA on the lipids selected by PLS-DA. Colored shapes indicate treatment groups as follows: red triangle, ambient pH:ambient DO; green cross, ambient pH:low DO; blue X, low pH:ambient DO; turquoise diamond, low pH:low DO.

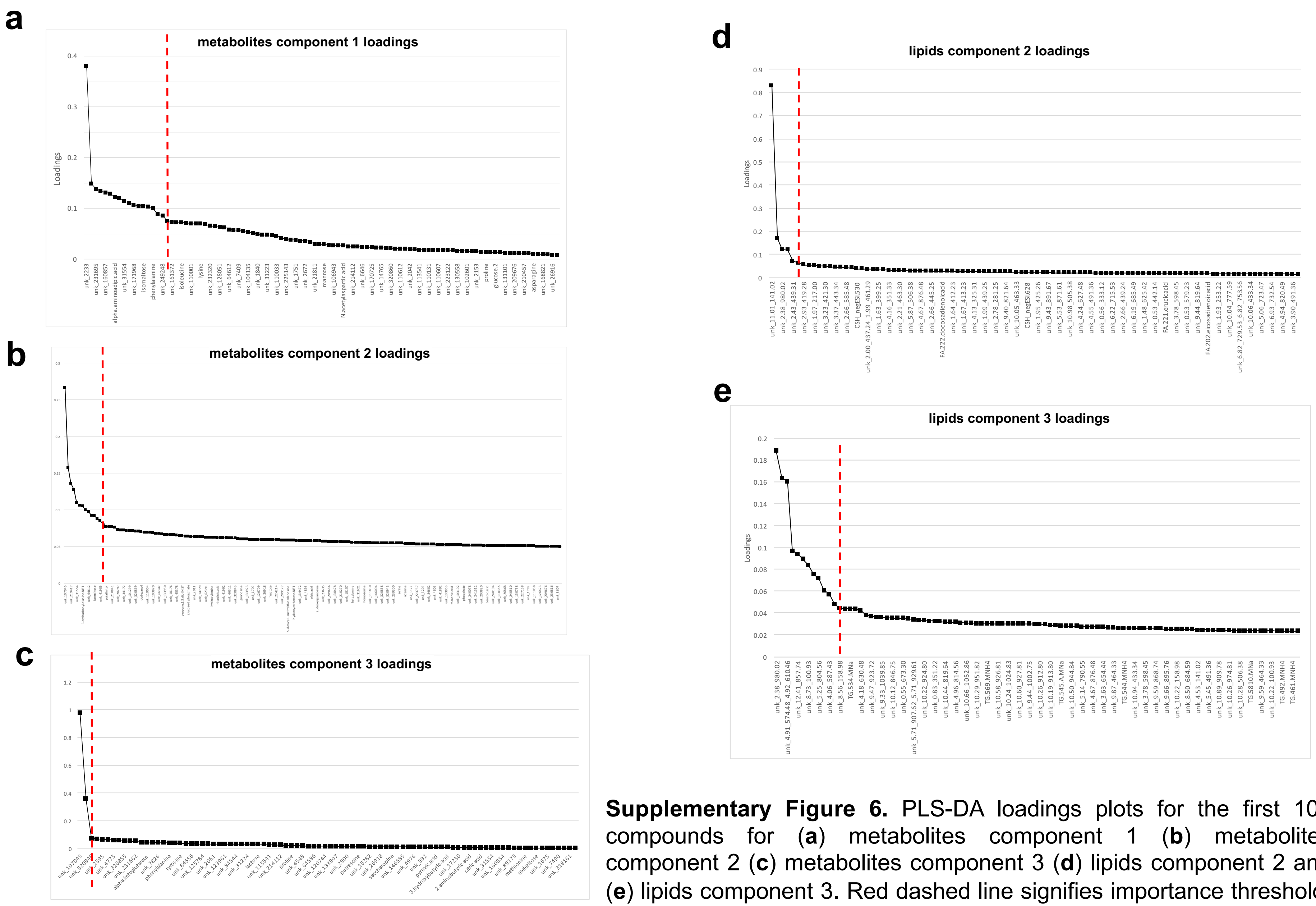


**Supplementary Figure 6.** PLS-DA loadings plots for the first 100 compounds for (**a**) metabolites component 1 (**b**) metabolites component 2 (**c**) metabolites component 3 (**d**) lipids component 2 and (**e**) lipids component 3. Red dashed line signifies importance threshold. X-axis labels are an abbreviated list of the 100 compounds. **Supplementary Table 7** contains PLS-DA loadings data for all compounds.

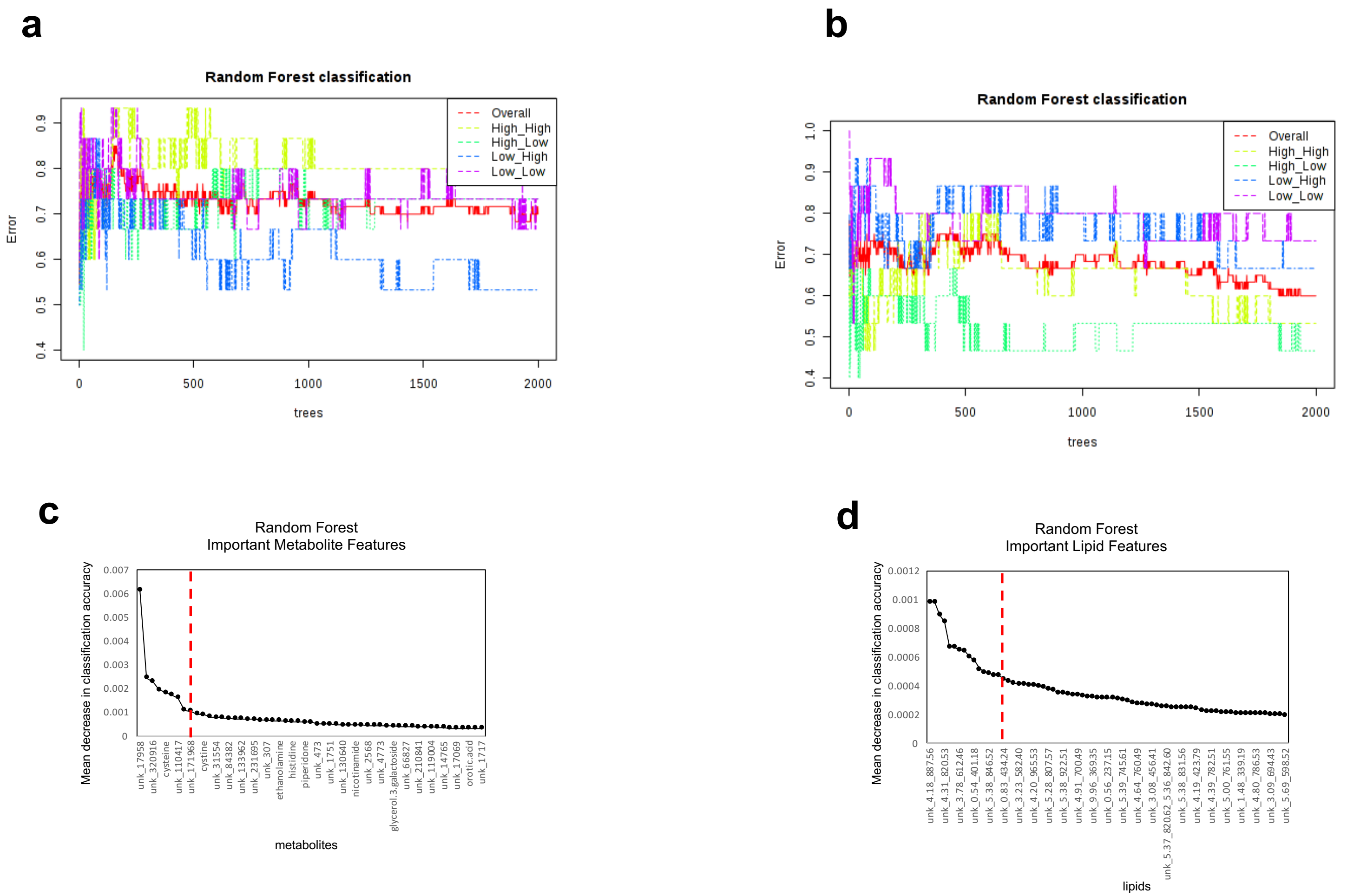


**Supplementary Figure 7.** Performance of the random forest classifier applied to the whole (**a**) metabolite and (**b**) lipid datasets. Random forest decrease in accuracy plot for (**c**) metabolite and (**d**) lipid data. Red line indicates importance threshold. X-axis labels are an abbreviated list of compounds. **Supplementary Table 8** contains mean decrease in random forest classification accuracy data for all compounds.

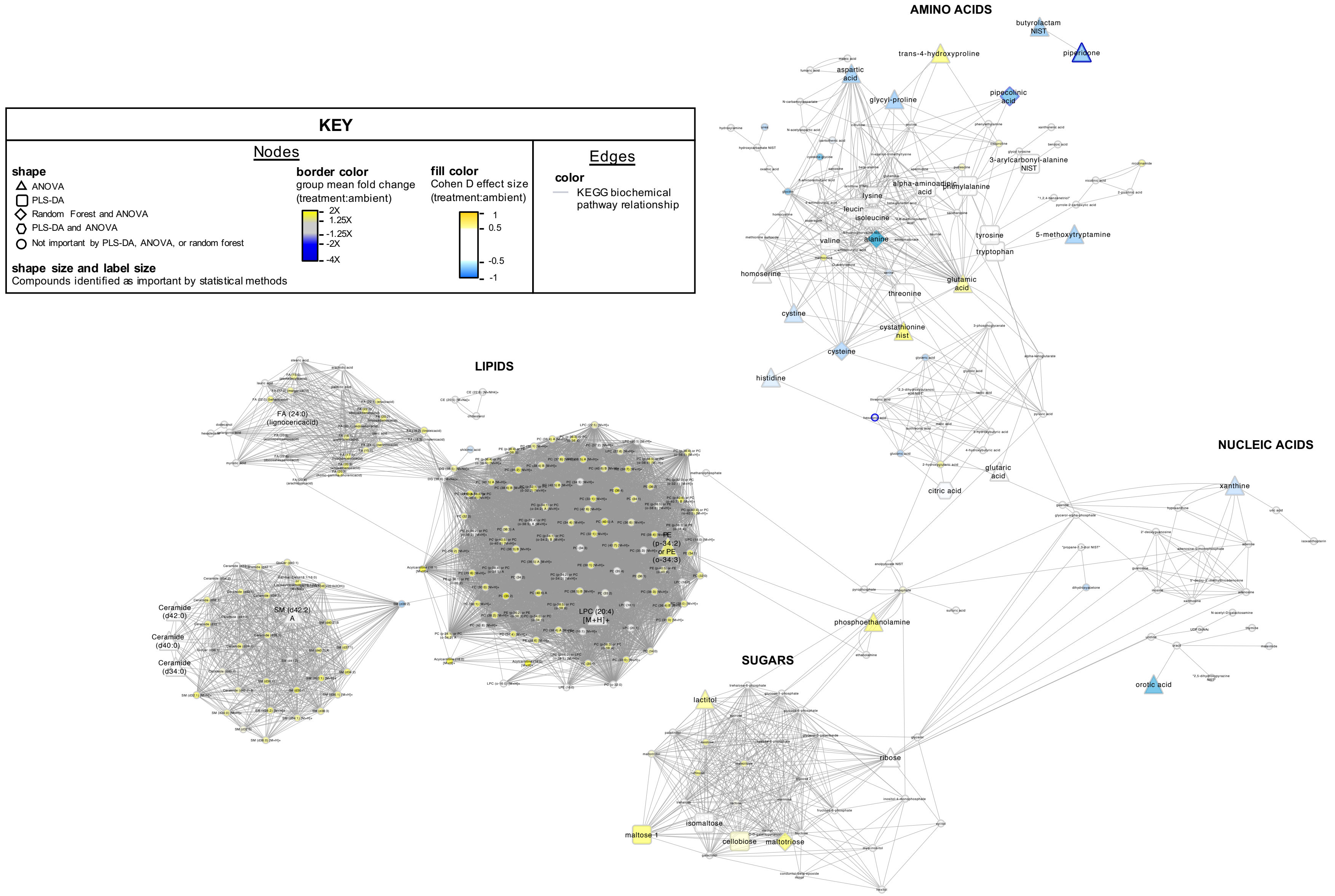


**Supplementary Figure 8.** Pathway analysis of compounds affected by low DO. Compounds are clustered by chemical similarity. Node fill color is colored by Cohen D effect size comparing the low DO treatment group to the ambient pH,ambient DO group, and node borders are colored by the low DO treatment group mean fold change relative to the ambient pH,ambient DO group. Node shapes indicate the statistical method(s) from which the compound was classified as important. Node shape and label size are enlarged if the compound was identified as important by a statistical method. Gray edges indicate nodes sharing a KEGG biochemical pathway(s).

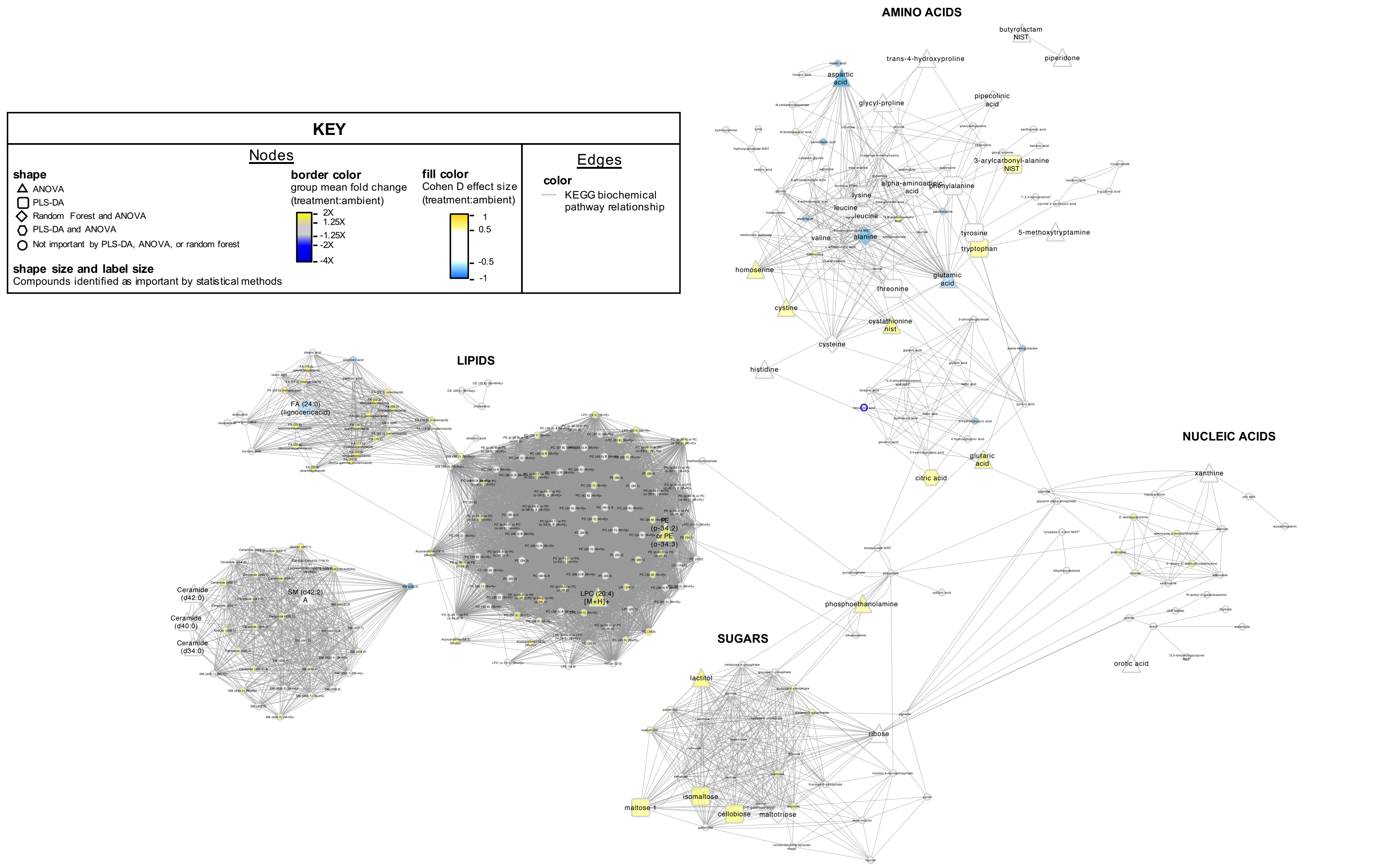


**Supplementary Figure 9.** Pathway analysis of compounds affected by low pH. Compounds are clustered by chemical similarity. Node fill color is colored by Cohen D effect size comparing the low pH treatment group to the ambient pH,ambient DO group, and node borders are colored by the low pH treatment group mean fold change relative to the ambient pH,ambient DO group. Node shapes indicate the statistical method(s) from which the compound was classified as important. Node shape and label size are enlarged if the compound was identified as important by a statistical method. Gray edges indicate nodes sharing a KEGG biochemical pathway(s).

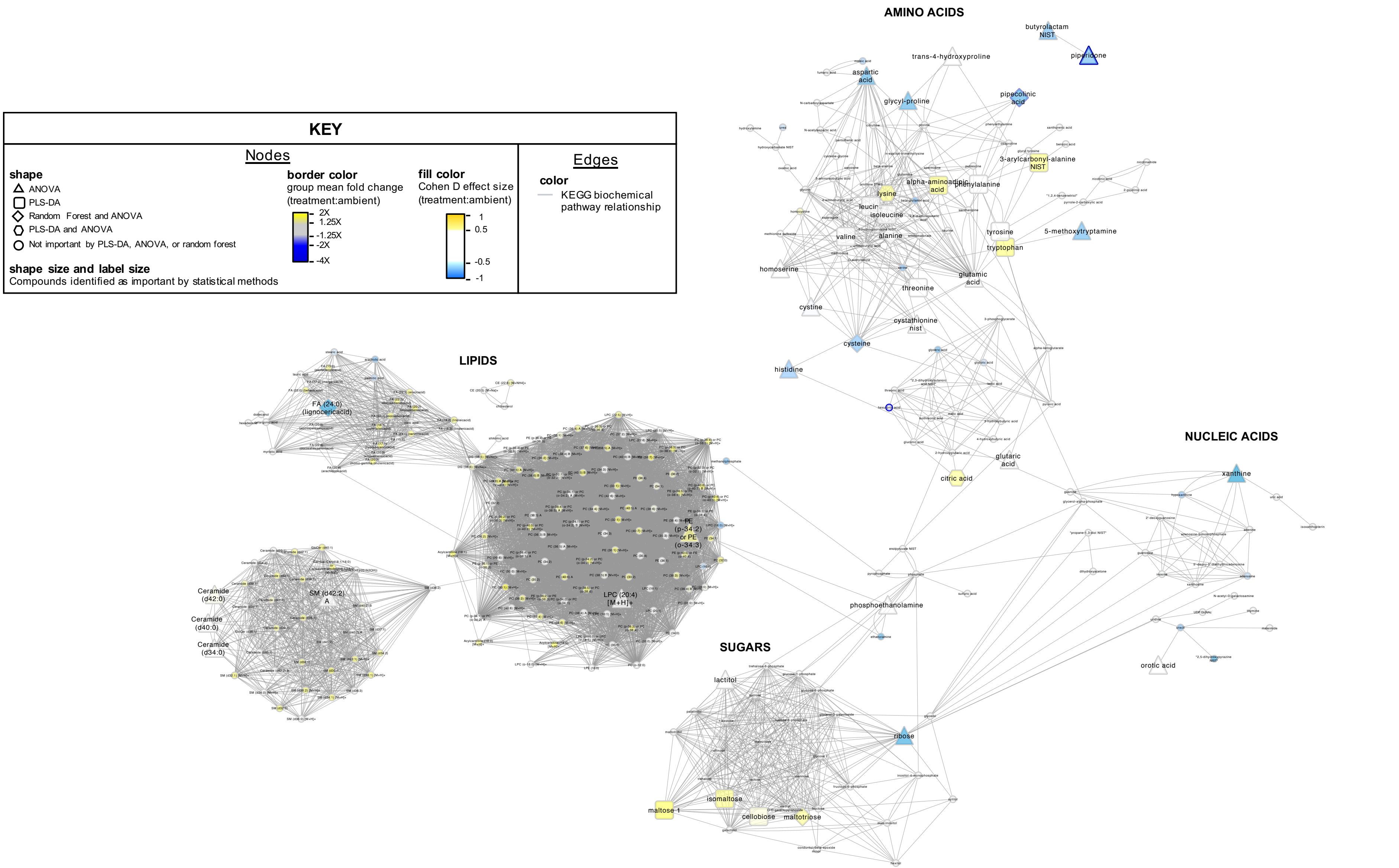


**Supplementary Figure 10.** Pathway analysis of compounds affected by combined low pH and low DO. Compounds are clustered by chemical similarity. Node fill color is colored by Cohen D effect size comparing the low pH,low DO treatment group to the ambient pH,ambient DO group, and node borders are colored by the low pH,low DO treatment group mean fold change relative to the ambient pH,ambient DO group. Node shapes indicate the statistical method(s) from which the compound was classified as important. Node shape and label size are enlarged if the compound was identified as important by a statistical method. Gray edges indicate nodes sharing a KEGG biochemical pathway(s).
